# Regulation of EBNA1 Protein Stability by PLOD1 Lysine Hydroxylase

**DOI:** 10.1101/2022.03.29.486189

**Authors:** Jayaraju Dheekollu, Andreas Wiedmer, Samantha S. Soldan, Leonardo Josué Castro Muñoz, Hsin-Yao Tang, David W. Speicher, Paul M. Lieberman

**Affiliations:** The Wistar Institute, Philadelphia, PA 19104 USA

**Keywords:** Epstein-Barr Virus, EBV, PLOD, OriP, episome maintenance, protein stability, Procollagen-Lysine Hydroxylase

## Abstract

Epstein-Barr virus (EBV) is a ubiquitous human γ-herpesvirus that is causally associated with various malignancies and autoimmune disease. Epstein-Barr Nuclear Antigen 1 (EBNA1) is the viral-encoded DNA binding protein required for viral episome maintenance and DNA replication during latent infection in proliferating cells. EBNA1 is known to be a highly stable protein, but its mechanism of protein stability is not completely understood. Proteomic analysis of EBNA1 revealed interaction with Procollagen Lysine-2 Oxoglutarate 5 Dioxygenase (PLOD) family of proteins. Depletion of PLOD1 by shRNA or inhibition with small molecule inhibitors 2,-2’ dipyridyl resulted in the loss of EBNA1 protein levels, along with a selective growth inhibition of EBV-positive lymphoid cells. PLOD1 depletion also caused a loss of EBV episomes from latently infected cells and inhibited *oriP*-dependent DNA replication. We used mass spectrometry to identify EBNA1 peptides with lysine hydroxylation at K460 or K461. Mutation of K460 to alanine or arginine abrogates EBNA1-driven DNA replication of *oriP, while K461 mutations enhanced replication*. These findings suggest that PLOD1 is a novel post-translational regulator of EBNA1 protein stability and function in viral plasmid replication, episome maintenance and host cell survival.

**Importance:** EBNA1 is essential for EBV latent infection and implicated in viral pathogenesis. We found that EBNA1 interacts with PLOD family of lysine hydroxylases and that this interaction is required for EBNA1 protein stability and function in viral persistence during viral latent infection. Identification of PLOD1 regulation of EBNA1 protein stability provide new opportunity to target EBNA1 for degradation in EBV associated disease.

## Introduction

Epstein-Barr Virus (EBV) is a human gammaherpesvirus that establishes life-long latent infection in over 90% of the adult population world-wide [1, 2]. EBV latent infection is a causal agent for several cancers, including Burkitt Lymphoma (BL), Nasopharyngeal Carcinoma (NPC), and post-transplant lymphoproliferative diseases (PTLD) [3-5]. EBV is also associated with several autoimmune diseases, especially multiple sclerosis (MS) where viral proteins have been implicated as the molecular mimic and trigger for auto-reactive antibodies and T-cells [6, 7].

Epstein-Barr Nuclear Antigen 1 (EBNA1) is the viral-encoded sequence-specific DNA-binding protein that binds to tandem repeats in the viral origin of plasmid replication (oriP) and is required for viral episome maintenance and plasmid replication during latent infection in proliferating cells [8, 9]. EBNA1 can also modulate transcription of viral and host genes, and interacts with host proteins that are implicated in viral oncogenesis, such as USP7 and CK2 [10-12]. EBNA1 is predominantly localized to the nucleus of infected cells, and is the most consistently detected protein in EBV-associated tumors. EBNA1 is also known to have a relatively long half-life (∼20 hrs) in B-cells [13]. EBNA1 stabilization is partly dependent on a central gly-ala repeat that resists proteolysis associated with MHC peptide presentation [14, 15]. However, EBNA1 interaction with other proteins and post-translational modifications may also contribute to its stability[16].

The Procollagen-Lysine,2-Oxoglutarate 5-Dioxygenases (PLODs) are required for the post-translational modification that allows collagen cross-links and maturation of extracellular matrix (reviewed in [17]. PLOD1, 2 and 3 have different roles in collagen modification including a glycosylase activity unique to PLOD3. PLODs are expressed at different levels in different tissue types. While inherited mutations in PLODs cause connective tissue disorders, such as Ehlers-Danlos syndrome [18], upregulation of PLODs have been associated with several cancers, including gastric cancers and hepatocellular carcinomas [17, 19-23]. A recent study has found that PLOD1 and 3 can interact with EBNA1 in AGS gastric cells with preferential binding to EBNA1 isoforms found in epithelial cancers [24]. Here, we further advance these pioneering studies to show that EBNA1 can interact with all three PLODs and that depletion of PLOD1, or small molecule inhibition of PLOD enzymatic activity leads to a loss of EBNA1 protein stability and function in *oriP*-dependent DNA replication and episome maintenance. We also provide evidence that EBNA1 is subject to lysine hydroxylation that regulates EBNA1 replication function at *oriP*.

## Results

### EBNA1 proteomics identifies interaction with PLOD family of lysine hydroxylase

We have previously reported an LC-MS/MS proteomic analysis of EBNA1 [25]. For these studies, FLAG-EBNA1 was expressed from stable *oriP-*containing episomes to enrich for cellular proteins that bound to EBNA1 in the functional context of the *oriP*. We report here the identification of PLOD1, 2, and 3 as proteins highly enriched in FLAG-EBNA1 fraction relative to the FLAG-vector control (**Fig. 1A and B**). We also identified the USP7, which has been well-characterized for its interaction with EBNA1, and P4HA2, a proline hydroxylase related to PLODs (**Fig. 1B**). RNA analysis of PLODs revealed that two isoforms of PLOD1 (A and B) were expressed at higher levels than PLOD2 or PLOD3 in EBV+ B-cell lines (**Supplementary Fig. S1**). We therefore focused our efforts on characterization of PLOD1 with EBNA1 in these B-lymphocytes. Immunoprecipitation (IP) with endogenous EBNA1 in Raji and Mutu I Burkitt lymphoma cell lines revealed selective enrichment of PLOD1 relative to IgG control (**Fig. 1C**). Similarly, reverse IP with PLOD1 in Raji and Mutu I cells revealed selective enrichment of EBNA1 relative to IgG control (**Fig. 1D**). Interestingly, EBNA1 species precipitated in PLOD1 IP had an additional EBNA1 reactive species (*) of slower mobility, suggesting potential EBNA1 post-translational modification when complexed with PLOD1.

**Figure 1.**
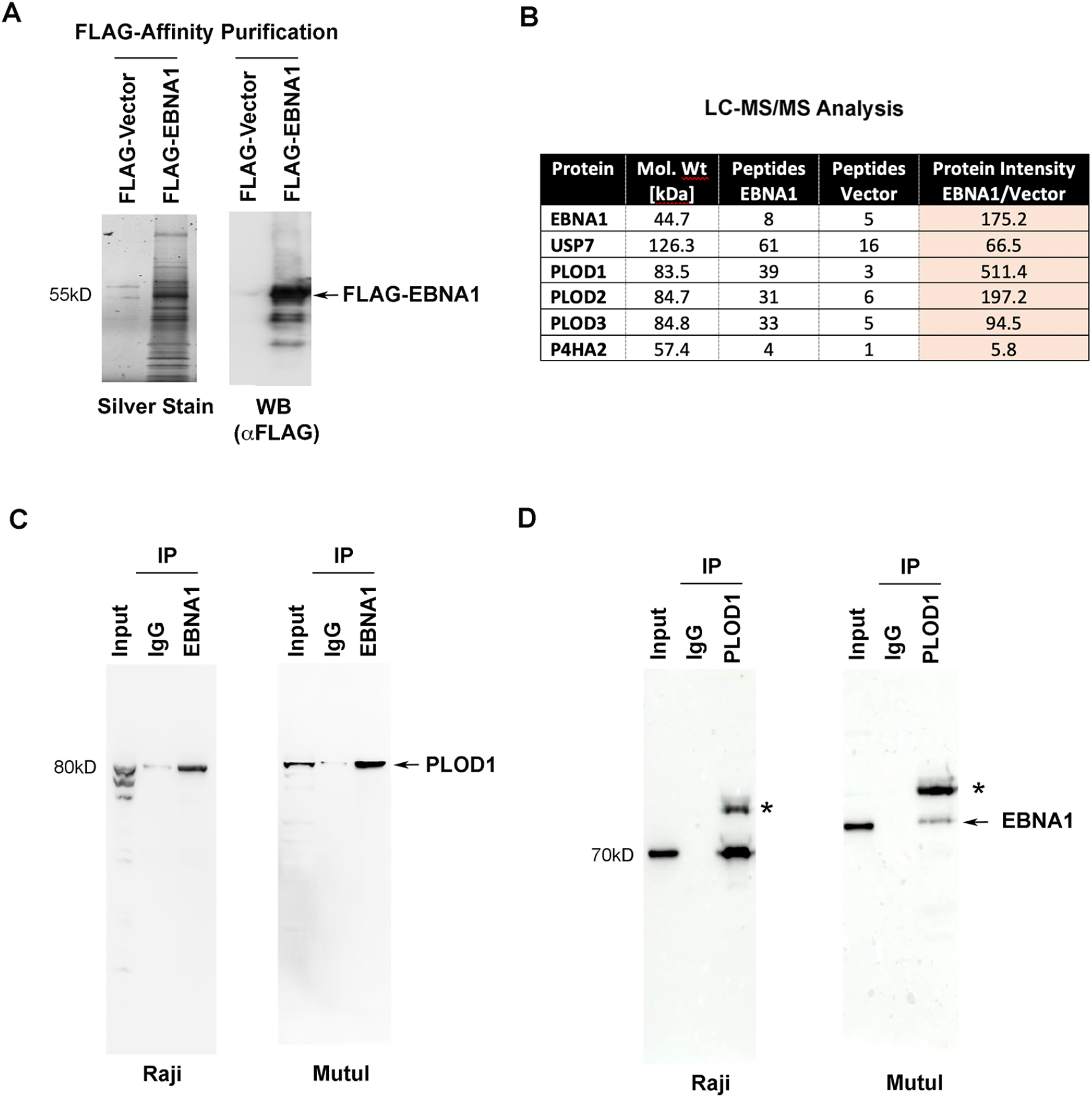
Identification of PLODs as EBNA1 interaction partners. **A)** FLAG-affinity purified proteins from HEK293T cells with stable expression of FLAG-EBNA1 or FLAG-Vector from *oriP* plasmids were analyzed by silver stain (left) or FLAG Western blot (WB, right). **B)** LC-MS/MS analysis of FLAG-EBNA1 associated proteins highlighting numbers of peptides identified for USP7, PLOD1, PLOD2, PLOD3, and P4HA2. **C)** Immunoprecipitation (IP) with EBNA1 or IgG control antibody from Raji (left) or MutuI (right) total cell extracts probed by Western blot with antibody to PLOD1. **D)** Same as in C, except reciprocal IP with PLOD1 antibody probed with antibody to EBNA1. * indicates a slower mobility form of EBNA1.

### Inhibitor of PLOD1 leads to loss of EBNA1

To investigate the potential effects of PLOD1 on EBNA1 protein expression, we first generated lentivirus expressing shRNA targeting PLOD1. We found that shRNA depletion of PLOD1 in Raji BL cells led to a significant loss of expression of PLOD1, indicating that shRNA knock-down was working efficiently (**Fig. 2A, top panel**). In the same knock-down of PLOD1, we observed a reduction in EBNA1 protein, along with a down-shift in EBNA1 mobility in SDS-PAGE Western blot (**Fig. 2A**). We also observed a similar change in EBNA2 and to a lesser extent that of LMP1, while cellular actin was not affected (**Fig. 2A**). To determine if these effects of PLOD1 protein depletion correlated with loss of PLOD1 enzymatic activity, we assayed the effects of a small molecule inhibitor of PLOD1. Bipyridine (also known as 2,2 dipyridil and referred to here as 2-DP) has been reported to have selective inhibition of PLOD1 [26]. We found that treatment of Raji and LCLs with 2-DP (100 µM) led to a loss of EBNA1 and EBNA2 in both cell types, with less of an effect on LMP1 or cellular actin (**Fig. 2B**), thus phenocopying shRNA depletion of PLOD1. To determine if the loss of EBNA1 protein levels were partly due to proteosome degradation, we assayed the effects of 2-DP in combination with proteosome inhibitor MG132 (**Fig. 2C**). We found that MG132 stabilized EBNA1 protein in the presence of 2-DP, suggesting that 2-DP leads to proteosomal degradation of EBNA1. EBNA1 protein can be destabilized by other small molecules, such as the HSP90 inhibitor 17-DMGA [27]. We found that 17-DMGA did not lead to the degradation of EBNA1 as did 2-DP under these conditions. Since 2-DP has the potential to chelate iron and induce hypoxia stress response, we compared the effects of 2-DP to treatment of CoCl_2_ a known inducer of hypoxic stress response through stabilization of HIF1A (**Fig. 2D**). We found that 2-DP led to a loss of PLOD1 and EBNA1 in both Raji and LCL, and stabilized HIF1A modestly in LCLs only. In contrast, CoCl2 stabilized HIF1A in both Raji and LCL, and reduced PLOD1 and EBNA1 in LCL, but had only weak effects on PLOD1 and EBNA1 in Raji cells. These findings suggest that 2-DP may inhibit PLOD1 through mechanisms distinct from HSP90 inhibition or hypoxia stress response, although there may be some cell-type dependent overlaps with these pathways.

**Figure 2.**
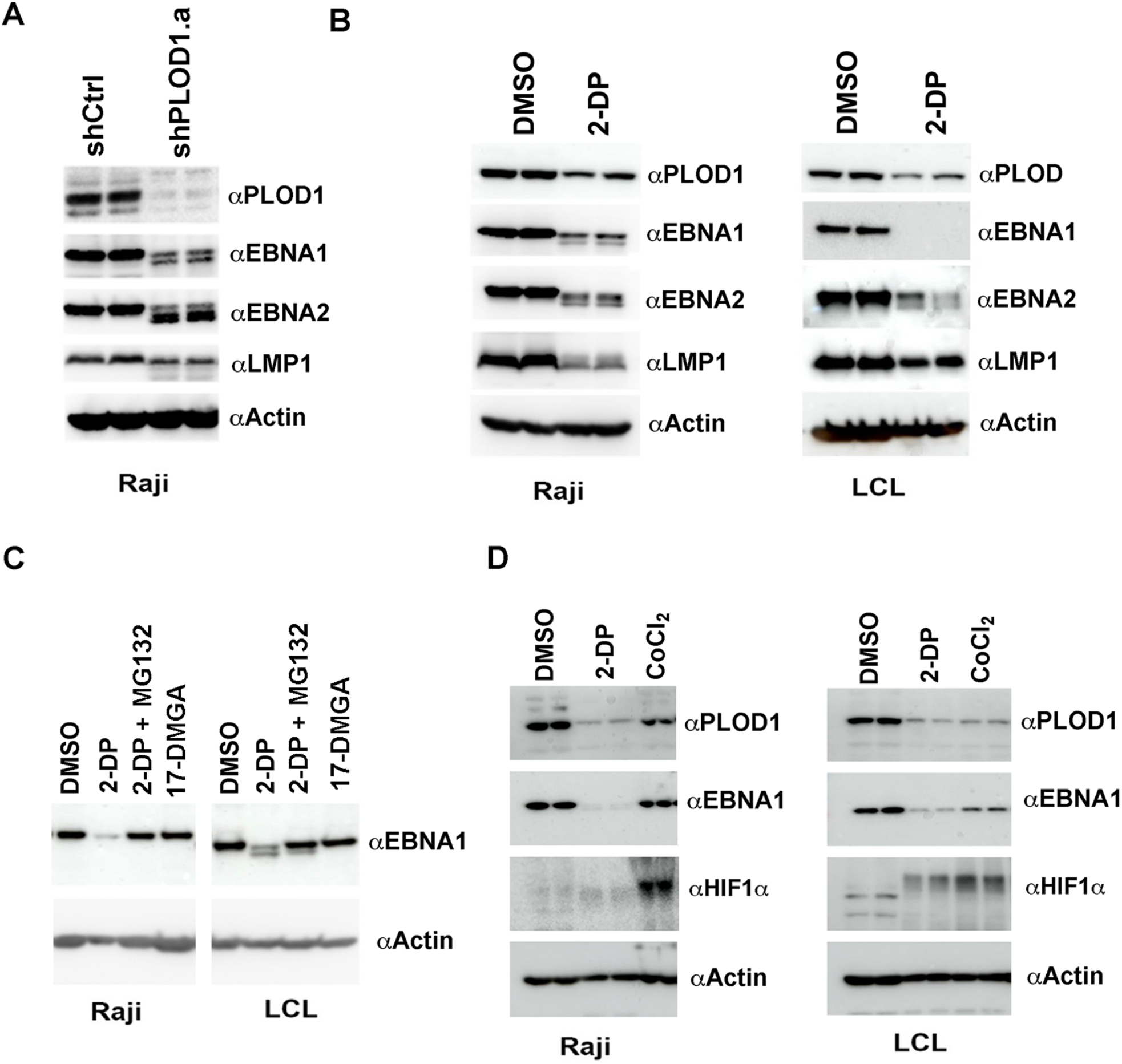
PLOD shRNA depletion and small molecule inhibition with 2-DP destabilize EBNA1 protein. **A)** Raji cells transduced with lentivirus shControl (shCtrl) or shPLOD1.a were assayed by Western blot for PLOD1, EBNA1, EBNA2, LMP1, or Actin. Each lane is a biological replicate. **B)** Raji (left) or LCL (right) treated with DMSO or 2,’2-dipyridil (2-DP, 200 µM) were assayed by Western blot for PLOD1, EBNA1, EBNA2, LMP1, or Actin. **C)** Raji (left) or LCL (right) were treated with DMSO, 2-DP (200 µM), 2-DP+MG132 (10 µM), or 17-DMGA (1 µM) for 48 hrs and assayed by Western lbot for EBNA1 (top) or Actin (bottom). **D)** Raji (left) or LCL (right) were treated with DMSO, 2-DP (200 µM), or CoCl_2_ (100 µM) for 48 hrs followed by Western blot for PLOD1, EBNA1, HIF1α, or Actin.

### Inhibition of PLOD1 selectively block EBV+ B cell survival

We next tested the effects of PLOD1 shRNA depletion and inhibition by 2-DP on EBV-dependent cell growth and survival. We compared EBV positive cells B-cell lines (Raji and MutuI BL and B95.8 transformed LCLs) with EBV negative B-lymphoma cell lines (BJAB and DG75). Cells were treated with 100 µM 2-DP for 2 days or with lentivirus transduction of shPLOD1 for 4 days followed by FACS profiling for propidium iodide (PI) and annexin V staining (**Fig. 3**). We found that both shPLOD1 and 2-DP induced a significant decrease in the percentage of proliferating/live cells (Q4) for EBV-positive MutuI, Raji, and LCL relative to EBV-negative BJAB and DG75. LCLs were particularly sensitive to shPLOD1-mediated depletion (**Fig.3B**). These findings suggest that EBV positive lymphoid cells are more sensitive than EBV negative lymphoid cells to loss of PLOD1 protein and its enzymatic activity.

**Figure 3.**
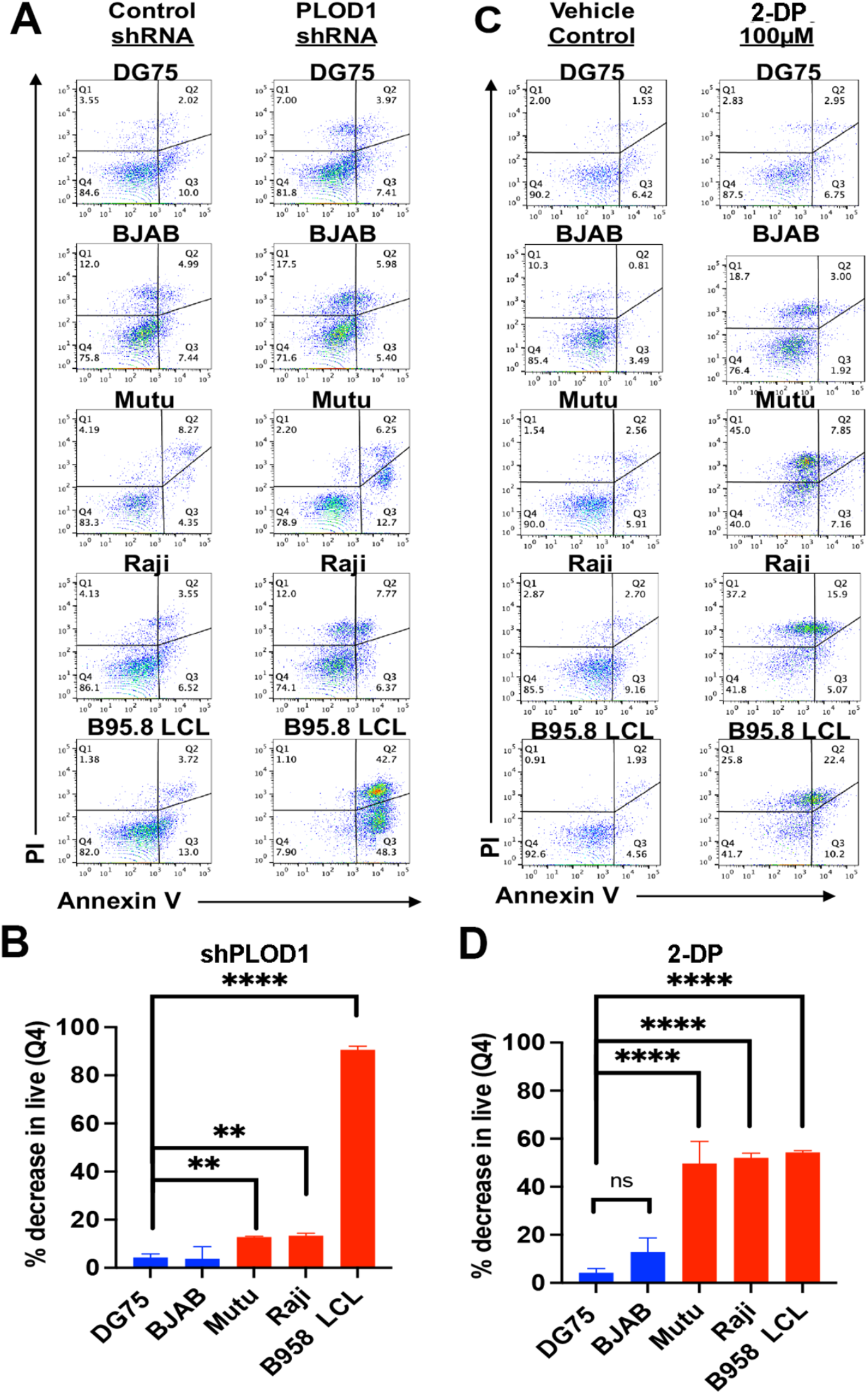
Selective inhibition of EBV-positive cells to PLOD1 depletion or inhibition by 2-DP. **A)** Flow cytometry analysis of cell viability using Propidium Iodide (PI, y-axis) and Annexin V (x-axis) for DG75, BJAB, MutuI, Raji, B95.8LCL transduced with control shRNA or PLOD1 shRNA. **B)** Quantification of the % decrease in live cells in treatments described in panel A. EBV positive cells (red bars), EBV negative cells (blue bars). **C)** Same as in panel A, except treatment with DMSO or 2-DP (200 µM). **D)** Quantification of the % decrease in live cells in treatments described in panel C. P-values determined by ordinary one-way ANOVA and Dunnett’s multiple comparison test ****<0.0001, **<0.01, *=0.0108.

### PLOD1 contributes to EBV episome maintenance in latently infected B-lymphocytes

We next assayed the effects of PLOD1 depletion on the maintenance of EBV episomes in two different BL (Mutu I and Raji) and LCL (transformed with B95-8 or Mutu virus) cell lines (**Fig. 4**). PFGE analysis revealed that shPLOD1 depletion caused a significant loss of EBV episomal DNA in each cell type (**Fig. 4A and B**). The efficiency of shPLOD1 depletion was measured by RT-qPCR and Western blot for each cell type (**Supplementary Fig S2**). EBV episome loss was striking despite relatively weak depletion of PLOD1 protein at this early time point prior to loss of cell viability.

**Figure 4.**
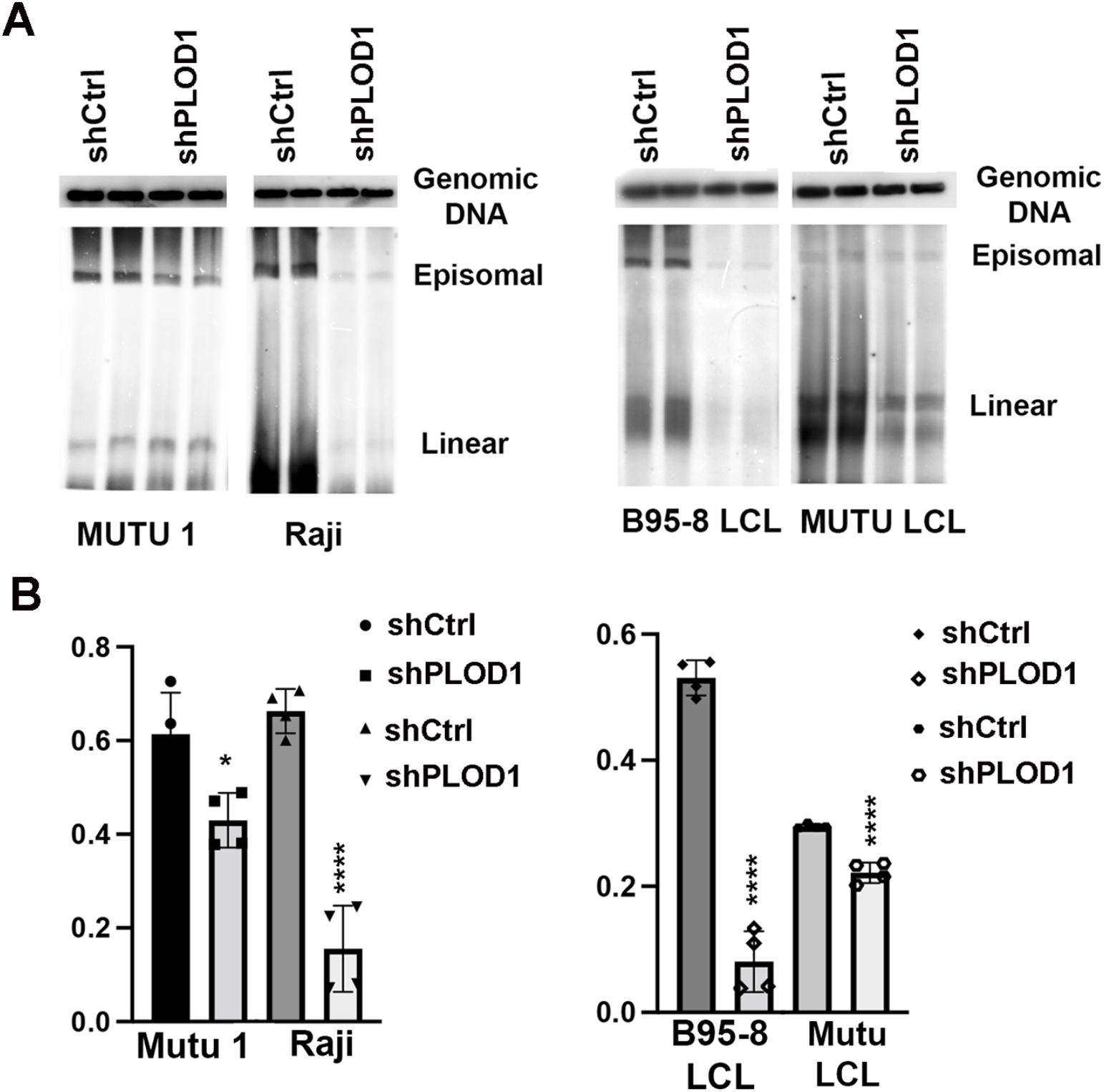
PLOD depletion causes loss of EBV episomes. **A)** PFGE analysis of MutuI, Raji, or B95-8 or Mutu LCLs transduced with lentivirus expressing shCtrl or shPLOD1 and analyzed by Southern blot for Genomic DNA (top) or EBV BamHI W repeat (lower panel) indicating viral episomes or linear genomes. **B)** Quantification EBV episomes from PFGE shown in panel C. P-values determined by ordinary one-way ANOVA and Dunnett’s multiple comparison test ****<0.0001, *=0.0108.

### PLOD1 contributes to EBNA1-dependent DNA replication

To determine if PLOD1 affected EBNA1 DNA replication function, we assayed transient plasmid replication in HEK293 cells transfected with *oriP*-containing plasmids that also expressed FLAG-EBNA1. We also assayed two different PLOD1 shRNAs, shPLOD1.a and shPLOD1.b for their ability to efficiently deplete PLOD1. While shPLOD1.a and shPLOD1.b led to a modest reduction in PLOD1 protein at this time point, the depletion on FLAG-EBNA1 expression was substantial (**Fig. 5A**). We then assayed the effect of shPLOD1 on EBNA1-dependent DNA replication. We found that both shPLOD1.a and shPLOD1.b substantially reduced *oriP*-dependent DNA replication, as measured by DpnI resistance assay and Southern blot detection of *oriP*-containing plasmid DNA (**Fig. 5B and C**). These findings further support the role for PLOD1 in the stabilization of EBNA1 protein levels, and its functional importance in for *oriP*-dependent DNA replication.

**Figure 5.**
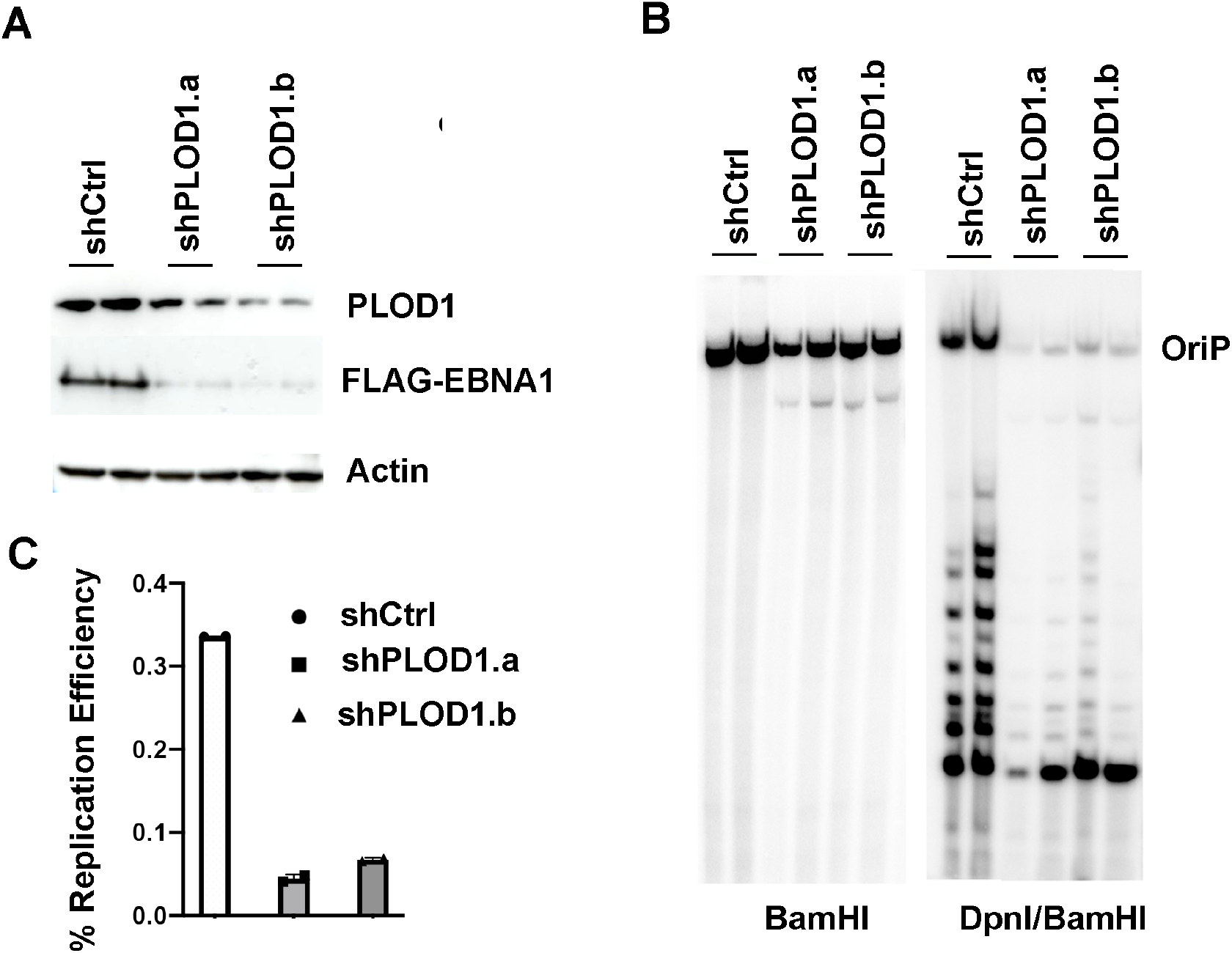
PLOD depletion inhibits EBNA1-dependent DNA replication of oriP plasmids. **A)** HEK293T cells transfected with oriP plasmids expressing FLAG-EBNA1 were transfected with expression vectors for shCtrl, shPLOD1.a, shPLOD1.b and assayed by Western blot for PLOD1 (top), FLAG-EBNA1 (middle) or Actin (bottom). **B)** oriP-plasmid replication for cells treated as in panel A assayed by Southern blot after BamHI digest (left) or DpnI/BamHI (right). Undigested linear oriP plasmid DNA is indicated. Each lane represents a biological duplicate. **C)** Quantification of % replicated oriP DNA for experiments shown in panel B.

### Lysine hydroxylation of EBNA1

To investigate the possibility that EBNA1 may be subject to post-translational modification through lysine hydroxylation, we performed LC-MS/MS analysis of immunoprecipitated EBNA1. We identified one peptide with a mass/charge (m/z) shift consistent with a single lysine hydroxylation (**Fig. 6A-C**). The EBNA1 peptide aa 416-465 had two potential lysine residues that could be hydroxylated, K460 and K461. PLOD1 typically hydroxylates lysines that precede glycine. We therefore first tested whether mutations in K461 impacted EBNA1 function in *oriP*-dependent DNA replication (**Fig. 6D-F**). We found that K461A had a modest stimulatory effect, while K461R had no significant effect on *oriP* replication (**Fig. 6D-F**). We next asked whether mutations in the neighboring K460 had any effects on oriP-DNA replication (**Fig. 6G-I**). We also included a mutation in K83A, which also has a PLOD1 consensus recognition site, and has been previously implicated in the PLOD1 interaction with the EBNA1 N-terminus. All EBNA1 mutants were expressed at similar levels in HEK293T cells (**Fig. 6G**). We found that K83A had a modest enhancement of *oriP* replication, while both K460A and K460R reduced oriP replication >5-fold (**Fig. 6H and I**). We also found that mutations in both K460A and K461A bound to *oriP* similar to wild-type EBNA1 as measured by ChIP assay, suggesting that these effects are not due to the disruption of EBNA1-DNA binding (**Supplementary Figs S3 and S4**). Taken together, these findings indicate that EBNA1 can be hydroxylated on either K460 or K461, and that mutations in K460 reduces EBNA1 replication function, but not its ability to bind at *oriP*.

**Figure 6.**
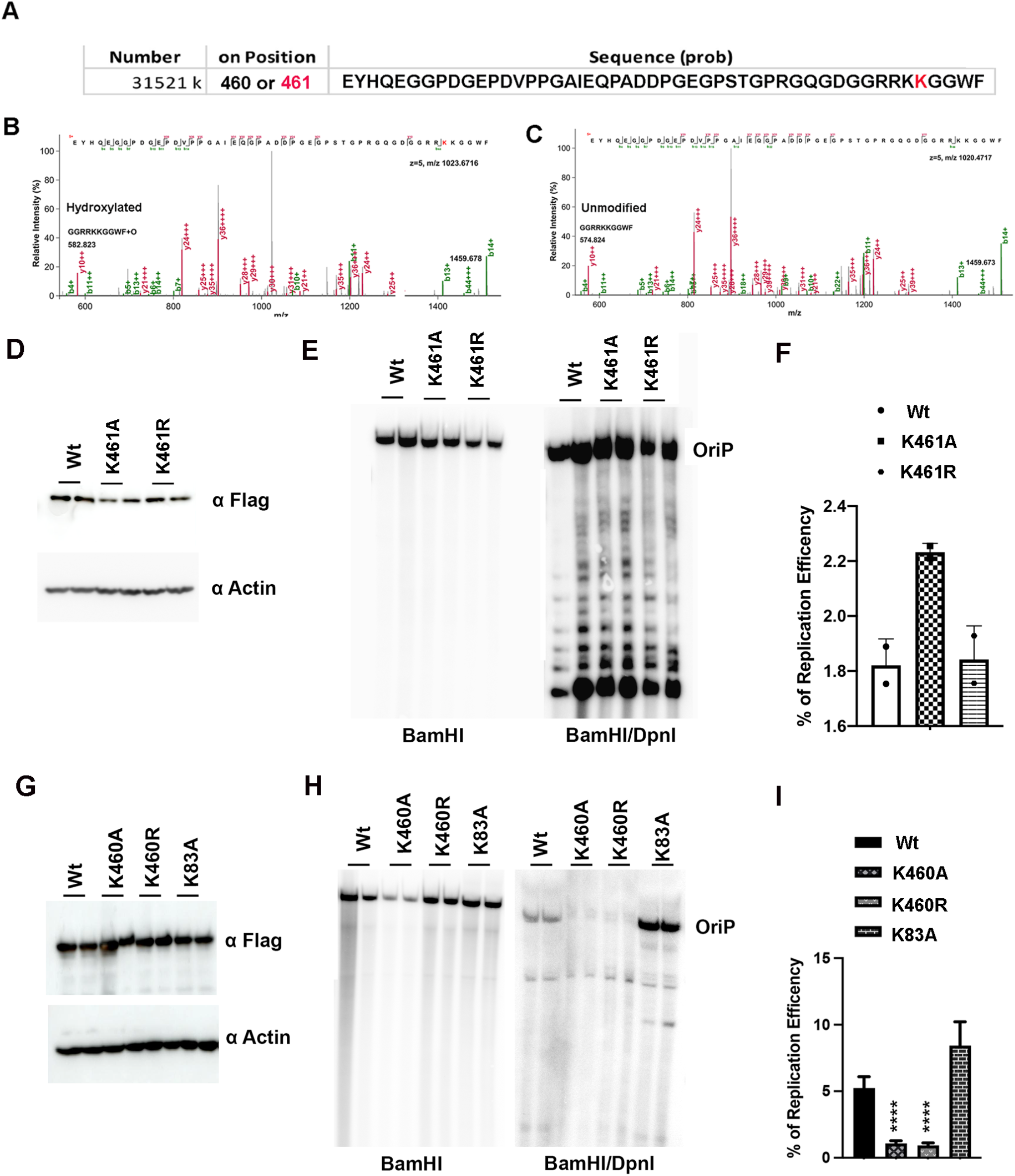
Evidence for lysine hydroxylation of EBNA1. **A)** Mass spectrometry (MS) of EBNA1 peptide with a mass/charge shift consistent with lysine hydroxylation. Consensus PLOD1 substrate recognition site at K461 highlighted in red. **B)** MS/MS spectrum of EBNA1 peptide with hydroxylation. **C)** MS/MS spectrum of the unmodified EBNA1 peptide. **D)** Western blot analysis of EBNA1 Wt, K461A, or K461R expressed in HEK293T cells. **E)** Southern blot analysis of oriP-dependent DNA replication for EBNA1 Wt, K461A, or K461R. **F)** Quantification of % oriP DNA replication shown in panel E. **G)** Western blot analysis of EBNA1 Wt, K460A, or K460R expressed in HEK293T cells. **H)** Southern blot analysis of *oriP*-dependent DNA replication for EBNA1 Wt, K460A, or K460R. **I)** Quantification of % *oriP* DNA replication shown in panel E. P-values determined by ordinary one-way ANOVA and Dunnett’s multiple comparison test ****<0.0001

## Discussion

EBNA1 is thought to be a highly stable protein in the nucleus of cells latently infected with EBV. Herein, we describe an EBNA1 interaction partner, PLOD1, that contributes to EBNA1 protein stability and essential functions in episome maintenance and DNA replication. We identified PLODs 1, 2, and 3 as EBNA1-associated proteins by LC-MS/MS and validated the interaction with PLOD1 antibody and coIP experiments in transfected 293HEK, and with native proteins in latently infected LCLs and BL cells. We found that shRNA depletion of PLOD1 led to a loss of EBNA1 protein levels in various B-cell types tested. PLOD1 depletion also led to a loss of EBNA2, suggesting that it may have more EBV substrates than just EBNA1. A small molecule inhibitor of PLOD1, namely 2-DP, phenocopies the effects of PLOD1 depletion. 2-DP could also induce HIF1alpha in some cell types, but CoCl_2_ induced hypoxia did not result in the same loss of EBNA1 protein stability. 2-DP and PLOD1 depletion led to loss of cell viability in an EBV-dependent manner. PLOD1 depletion led to a loss of EBV episomes in BL and LCL cells, as well as a loss of *oriP*-dependent DNA replication in HEK293 cells. Finally, we used mass spectrometry to identify EBNA1 peptides with mass/charge shifts consistent with lysine hydroxylation at K460 or K461. While mutations at K461 had only small effects on EBNA1 replication activity, mutations of K460 strongly attenuated EBNA1 replication function. We conclude that PLOD1 regulates EBNA1 protein stabilization and function in regulation of EBV latency, that EBNA1 is hydroxylated on lysine K460 or K461, and that mutations in K461 lead to a loss of EBNA1 replication function.

PLOD1 and PLOD3 have previously been reported to interact with EBNA1 [24]. In this earlier study, PLOD1 was found to bind preferentially to EBNA1 with a polymorphism (T85A) frequently associated with NPC and EBVaGC [24]. PLOD1 interaction with EBNA1 was found to be dependent on K83, a lysine residue in the N-terminal domain that also conforms to a consensus substrate for PLOD1 hydroxylation. EBNA1 N-terminal domain is involved in transcriptional activation, but it remains to be shown whether PLOD1 contributes to the transcriptional activation function of EBNA1. It was proposed that EBNA1 interaction with PLOD1 may sequester PLOD1 away from other substrates, such as procollagen, to drive tumorigenesis [24]. Our findings are mostly consistent with these previous findings, but provide new information on the role of PLODs in the regulation of EBNA1 protein stabilization and function in DNA replication and episome maintenance. Our data suggests that PLODs directly affect EBNA1 through post-translational modification and stabilization.

Overexpression of PLOD proteins have been implicated in several human cancers [17]. Our findings suggest that PLOD1 can both bind and regulate EBNA1 protein stability. PLOD1 depletion also led to large effects on EBV episome maintenance and *oriP* DNA replication, suggesting that PLOD1 binding or modification of EBNA1 may contribute to these activities beyond mere protein stabilization. Protein stabilization is integrally linked to many functions, including transcriptional activation [28] and replication origin function [29]. Protein hydroxylation of proline regulates HIF1A in the hypoxic response [30] and PLODs are well-characterized for modifying pro-collagen in the maturation of extracellular matrix [31]. PLODs utilize iron and alpha-ketoglutarate as cofactors, so it is likely that small molecules that alter these components, such as iron chelators, would have the potential to inhibit PLODs, as well as other iron-dependent enzymes. The precise role of PLODs in regulation of EBNA1 and potentially other EBV proteins, such as EBNA2, remain to be further investigated. Our findings suggest that a PLOD-dependent pathway is involved in maintaining EBNA1 stability and function, and that this may be exploited for disruption of EBV latency and treatment of EBV-associated disease.

## Materials and Methods

### Cells, Plasmids, and shRNAs

EBV-positive Burkitt’s lymphoma cells MutuI, Raji, MutuI virus-derived lymphoblastoid cell line (LCL) and, B95-8 LCL were grown in RPMI 1640 medium (Gibco BRL) containing 15% fetal bovine serum and antibiotics penicillin and streptomycin (50 U/ml). HEK 293T cells were culture in Dulbecco’s modified Eagle’s medium (DMEM) with 10% fetal bovine serum and antibiotics. All the cells were cultured at 37°C and 5% CO_2_ environment. Mammalian expression vector for Flag-EBNA1 contained B95-8 EBNA1 lacking the GA repeats (aa 101-324) under the control of CMV-3XFLAG promoter in a plasmid derived from pREP10 (Clontech) containing, *oriP*, GFP, and hygromycin resistance [25]. Small hairpin RNAs (shRNAs) for Plod1 (shPlod1), and the control (shControl) were obtained from the Sigma/TRC (The RNAi Consortium) collection of targeted shRNA plasmid library (TRC no. 62248, 62249, 62259, 62251and 62252). Lentivirus particles were generated in 293T-derived packaging cell lines.

### Drug Treatments

Raji and LCLs (2 × 10^5^ cells/ mL) were treated with vehicle control (DMSO; 0.016%, vol/vol) or 2-DP (200 μM) or 17-DMAG (1 μM) or CoCl_2_ (100 μM) for 48 hr. Cells were harvested and the Western blots were performed. For MG132 studies cells were treated with either vehicle control (DMSO; 0.016%, vol/vol) or 2-DP (200 μM) for 24 hr fallowed by adding MG132 to a final concentration of 10 μM and continue the treatment for another 24 hr.

### Site-directed mutagenesis

Primers were designed to generate the point mutations (K461A and K461R) in CMV Flag-EBNA1 containing oriP and hygromycin resistance plasmid (N2624). A two-stage PCR protocol for site-directed mutagenesis was adapted from Stratagene [25]. Following DpnI digestion and heat inactivation, PCR products were transformed into DH5α cells. Purified plasmids from colonies were sequenced to confirm the mutation.

### Western blots

The PVDF membranes were blotted with the following antibodies: anti-β-actin-peroxidase (Catalog NO. A3854; Sigma-Aldrich), anti-EBNA1 mouse monoclonal antibody (Catalog NO. sc-81581; Scbt), and anti-Flag M2-peroxidase (horseradish peroxidase [HRP]) (Sigma-Aldrich, cat no. A8592), anti-PLOD1 rabbit polyclonal (Catalog NO. HPA039137; Millipore), anti-PLOD1 rabbit polyclonal (Catalog NO.LS-C482920; LSBio), anti-EBNA2 rat polyclonal (Catalog NO. 50175912; Fisher), anti-LMP1 mouse monoclonal (Catalog NO. M0897; Dako), anti-EBNA1 rabbit polyclonal antibodies (custom prepared at Pocono Rabbit Farm), and imaging on a Amersham Imager 680.

### Chromatin Immunoprecipitation (ChIP)

ChIP assays were performed as previously described [32]. Briefly, 293T (∼1 × 10^6^ cells) were plated in 10 cm dishes. 24 h later cells were transfected with Lipofectamine 2000 (12 µl, Invitrogen) and 4 µg *oriP* plasmids expressing either FLAG-B95-8 EBNA 1or lysine mutation. Cells were split after 48 h, and then harvested at 72 h post transfection for ChIP assay. TaqMan qPCR performed using primers and probe designed by Themo-Fisher at *oriP*). Antibodies used were as follows: anti-IgG mouse monoclonal (Santa Cruz Biotechnology), anti-Flag resin (Catalog NO. M8823; Sigma-Aldrich)

### Plasmid replication assays

Plasmid DNA replication assays have been described previously [25, 33]. Briefly, 293T (∼1 × 10^6^ cells) were plated in 10 cm dishes. 24 h later cells were transfected with Lipofectamine 2000 (12 µl, Invitrogen) and 4 µg *oriP* plasmids expressing either FLAG-B95-8 EBNA 1, with shPLOD1 or shControl plasmids. Cells were split after 48 h, and then harvested at 72 h post transfection for both episomal DNA and protein. Episomal DNA was extracted by Hirt Lysis [34]. The DNA pellets were dissolved in 150 µl of 10 mM Tris HCl, 1 mM EDTA buffer (pH 7.6) and 15 µl was subjected to restriction digestion with BamHI alone and 135 µl was subjected to BamHI and DpnI digestion overnight at 37° C. DNA was extracted with phenol: chloroform (1:1), precipitated, and electrophoresed on a 0.9% agarose gel and transferred to a nylon membrane (PerkinElmer) for southern blotting. Blots were visualized and quantified using a Typhoon 9410

### shRNA-mediated knockdown of PLOD1

EBV-positive cells were infected by spin infection with lentivirus expressing shPLOD1, or shControl shRNA. At 48 h post-infection, 1.0 to 2.5 μg/ml puromycin was added to the media, and cell pools were selected for puromycin resistance. pLKO.1 vector-based shRNA constructs for were generated with target sequence 5′-T -3′ (shPLOD1). shControl was generated in pLKO.1 vector with target sequence 5′-TTATCGCGCATATCACGCG-3′. Lentiviruses were produced by cotransfection with envelope and packaging vectors pMD2.G and pSPAX2 in 293T cells. MutuI, Raji, or LCL cells were infected with lentiviruses carrying pLKO.1-puro vectors by spin-infection at 450 g for 90 minutes at room temperature. The cell pellets were resuspended and incubated in fresh RPMI medium, then treated with 2.5 µg/ml puromycin at 48 hrs after the infection. The RPMI medium with 2.5 µg/ml puromycin was replaced every 2 to 3 days. The cells were collected after 7 days of puromycin selection, then subject to following assays.

### EBV episome maintenance by pulsed-field electrophoresis

MutuI, Raji, and LCLs were infected with lentivirus. After 5 days of puromycin selection, cells were resuspended in 1× phosphate-buffered saline (PBS) and an equal amount of 2% agarose to form agarose plugs containing 1 × 10^6^ cells that were then incubated for 48 h at 50°C in lysis buffer (0.2 M EDTA [pH 8.0], 1% sodium lauryl sulfate, 1 mg/ml proteinase K). The agarose plugs were washed twice in TE buffer (10 mM Tris [pH 7.5] and 1 mM EDTA). Pulsed-field gel electrophoresis (PFGE) was performed for 23 h at 14°C with an initial switch time of 60 s and a final switch time of 120 s at 6 V/cm and an included angle of 120° as described previously (Bio-Rad CHEF Mapper) [35]. DNA was transferred to nylon membranes by established methods for Southern blotting [36]. The DNA was then detected by hybridization with α-^32^P-labeled probe specific for the EBV WP region and visualized with a Typhoon 9410 variable-mode imager (GE Healthcare Life Sciences).

### Immunoprecipitation

Cells were extracted with lysis buffer (20 mM Tris-HCl [pH 7.4], 1 mM EDTA, 0.1 mM EGTA, 2 mM MgCl_2_, 150 mM NaCl, 1 mM Na_3_VO_4_, 1 mM NaF, 20 mM sodium glycerophosphate, 5% glycerol, 1% Triton X-100, 0.5% sodium dodecyl sulfate, 1× protease inhibitors [Sigma], 1× phosphatase inhibitors [Sigma], and 1 mM phenylmethylsulfonyl fluoride [PMSF]). After rotation for 60 min at 4°C, the lysate was centrifuged for 20 min at 16,000 × *g*, and the supernatant was recovered. The cleared extracts were used for immunoprecipitation with antibodies as indicated in the figures.

### RNA analysis

Total RNA was extracted from EBV positive cells using TRIzol (Ambion) and then further treated with DNase I (New England Biolabs). Two micrograms of total RNA were reverse transcribed using random decamers (Ambion) and Superscript IV RNase H^−^ reverse transcriptase (Invitrogen). Specific primer sets were used in real-time quantitative PCR (qPCR) assays to measure Plod1a, Plod1b, Plod2 and, Plod3 levels. The values for the relative levels were calculated by ΔΔCT method.

### Flag-EBNA1 purification

293T cells were transfected with pCMV-Flag-EBNA1 OriP or Flag Vector plasmids. The cells were collected after 10 days post-transfection and washed once in 1X PBS. Cells (∼10^8^) were lysed in 50 ml of Lysis buffer (50 mM Tris pH 7.5, 150 mM NaCl, 0.5% Nonidet P40, 0.5% SDS, 1 mM EDTA), 1 mM PMSF, Protease inhibitors (Catalog NO. P8340; Sigma-Aldrich) and Phosphatase inhibitors (Catalog NO. 4906837001; Roche). Lysate were spin at 16000 for 10 min and immunoprecipitated with 100 µl of Anti-Flag resin (Catalog NO. M8823; Sigma-Aldrich). Complexes were washed three times with lysis buffer containing 300 mM NaCl, 1 mM PMSF, Protease inhibitors (Catalog NO. P8340; Sigma-Aldrich) and Phosphatase inhibitors (Catalog NO. 4906837001; Roche), and eluted with Flag peptide.

For EBNA 1 bound protein identification, 30 mg of Flag EBNA 1 complexes were run on a 10% precast gel (Invitrogen) for 1.5 cm and the gel was Coomassie stained. The entire stained gel regions were excised and digested with trypsin. Liquid chromatography tandem mass spectrometry (LC-MS/MS) analysis was performed using a Q Exactive HF mass spectrometer (ThermoFisher Scientific) coupled with a Nano-ACQUITY UPLC system (Waters). Samples were injected onto a UPLC Symmetry trap column (180 μm i.d. x 2 cm packed with 5 μm C18 resin; Waters), and peptides were separated by reversed phase HPLC on a BEH C18 nanocapillary analytical column (75 μm i.d. x 25 cm, 1.7 μm particle size; Waters) using a 2-h gradient formed by solvent A (0.1% formic acid in water) and solvent B (0.1% formic acid in acetonitrile). Eluted peptides were analyzed by the mass spectrometer set to repetitively scan m/z from 400 to 2000 in positive ion mode. The full MS scan was collected at 60,000 resolution followed by data-dependent MS/MS scans at 15,000 resolution on the 20 most abundant ions exceeding a minimum threshold of 20,000. Peptide match was set as preferred, exclude isotope option and charge-state screening were enabled to reject unassigned and single charged ions. Peptide sequences were identified using MaxQuant 1.5.2.8 [37]. MS/MS spectra were searched against a UniProt human protein database, EBNA1 protein sequence and a common contaminants database using full tryptic specificity with up to two missed cleavages, static carbamidomethylation of Cys, variable oxidation of Met, and variable protein N-terminal acetylation. Consensus identification lists were generated with false discovery rates set at 1% for protein and peptide identifications. Fold change was calculated using the protein intensity values.

### Mass Spectrometry

To identify post-translation modifications of EBNA 1, Flag-EBNA 1 complexes were washed three times with buffer contains 500 mM NaCl then Flag-EBNA 1 was eluted with 3X flag peptide and electrophoresed into an SDS-gel for a short distance. Gel regions containing Flag-EBNA 1 were digested separately with trypsin and chymotrypsin. Digests were analyzed by LC-MS/MS as described above. The MS data were searched using MaxQuant 1.6.2.3 [37]. Modifications searched were static carbamidomethylation of Cys, and variable Met oxidation, lysine hydroxylation, proline hydroxylation and protein N-terminal acetylation. Consensus identification lists were generated with false discovery rates set at 1% for protein, peptide, and site identifications.

### Cell viability assays

Cell viability was assessed 72 hours after 2’2-dipyridyl treatment using Resazurin cell proliferation/viability assay. In brief, EBV positive and negative cells were seeded onto 96-well plates and cultured overnight, followed by treatment over a ten-point concentration range of two-fold dilutions of 2’2-dipyridyl (0.39mM, 0.781mM, 1.56mM, 3.12mM, 6.25mM, 12,5 mM, 25 mM, 50 mM, 100 mM, 200 mM) (Sigma) plated in quadruplicate wells in 200 μL RPMI 1640 medium supplemented with 10% fetal bovine serum for 72 hours. As positive and negative controls, DMSO alone (0.4%) and puromycin (20 µg/ml) treated wells, respectively, were also plated in quadruplicate wells. At the end of the treatment, 20 μL of 500 mM Resazurin solution was added to each well and incubated for 6 hours at 37°C. The absorbance of each well was then detected at 590 nm under a microplate reader (CLARIOstarPlus, BMG Labtech). Cell viability was calculated as the ratio of the absorbance value to that of the control group (%) treated with 20 mg /ml puromycin.

### Cell apoptosis assay with flow cytometry

Apoptotic cells were detected using the FITC Annexin V Apoptosis Detection Kit (cat# ab14085, Abcam). EBV positive and negative cells were infected with lentivirus shCtrl or shplod1. After 48 hours post infection puromycin was added and selection for 3 days. The cells were then stained with Annexin V-FITC and PI according to the manufacturer’s instructions and the LSR14 Flow Cytometer (BD Biosciences). Cells were identified as viable, dead, or early or late apoptotic cells, and the percent decrease in live cell population (Q4: Annexin V(-)PI(-) was calculated as [Q4 control-Q4 treated/Q4 control] × 100 under each experimental condition.

## Data Availability

All data is available in the manuscript or supplementary data files.

## Acknowledgements

We thank members of the Wistar Cancer Center Cores in Proteomics, Genomics, and Flow Cytometry for their excellent technical support. This work was supported by grants from NIH DE017336, AI53508, CA140652, CA093606 to PML, R50 CA221838 to HYT and P30 Cancer Center Support Grant P30 CA010815 to the Wistar Institute (D. Altieri).

## Supplementary Figures

**Supplementary Figure S1.**
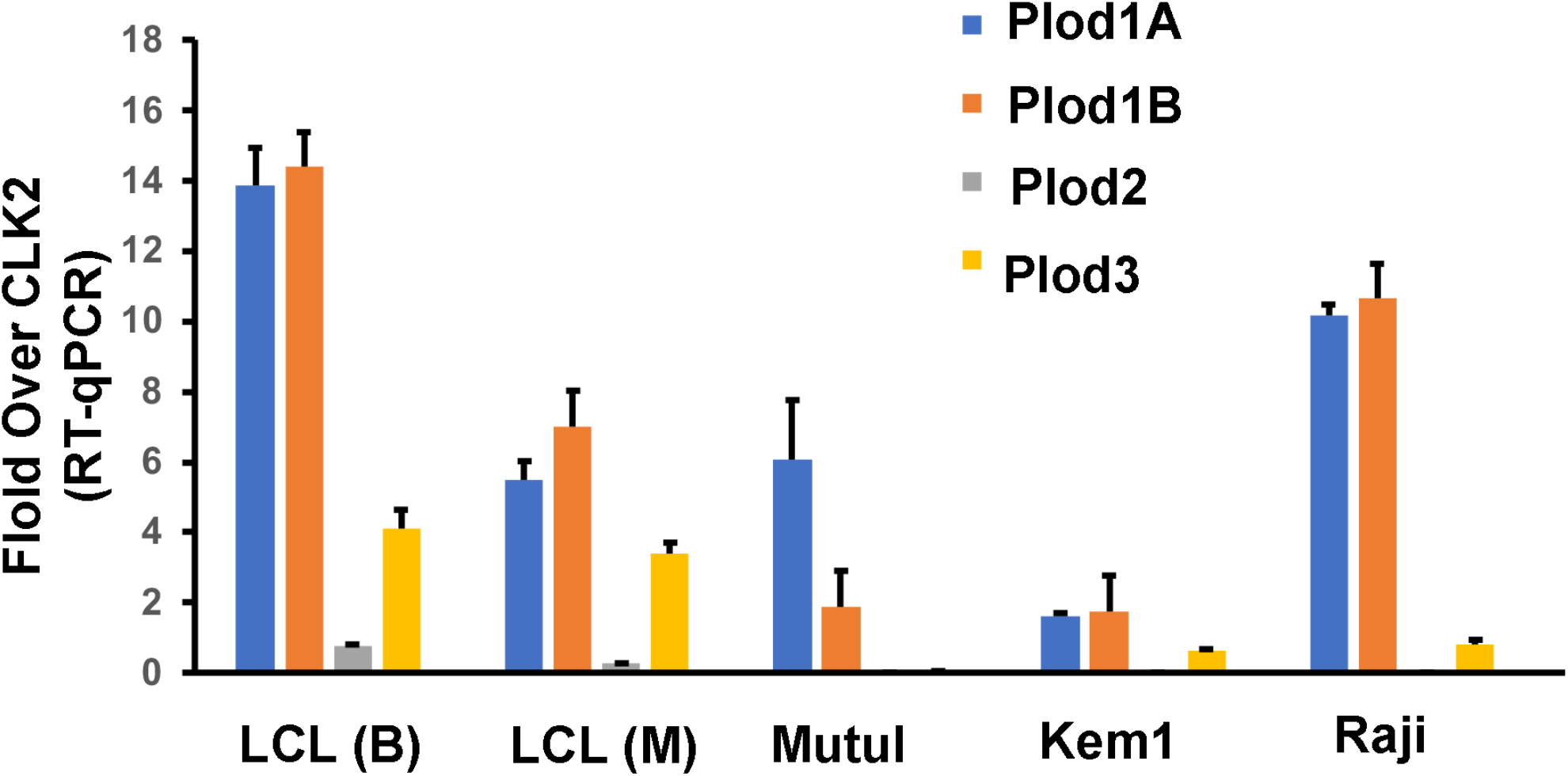
RNA expression of PLODs in lymphoid cells. RT-qPCR analysis of Plod1A, Plod1B, Plod2 and Plod3 transcripts in LCLs generated with B95.8 (B) or Mutu I (M) virus, or BL lines MutuI, Kem1, and Raji. Error bars are standard deviation, n=3 technical replicates.

**Supplementary Figure S2.**
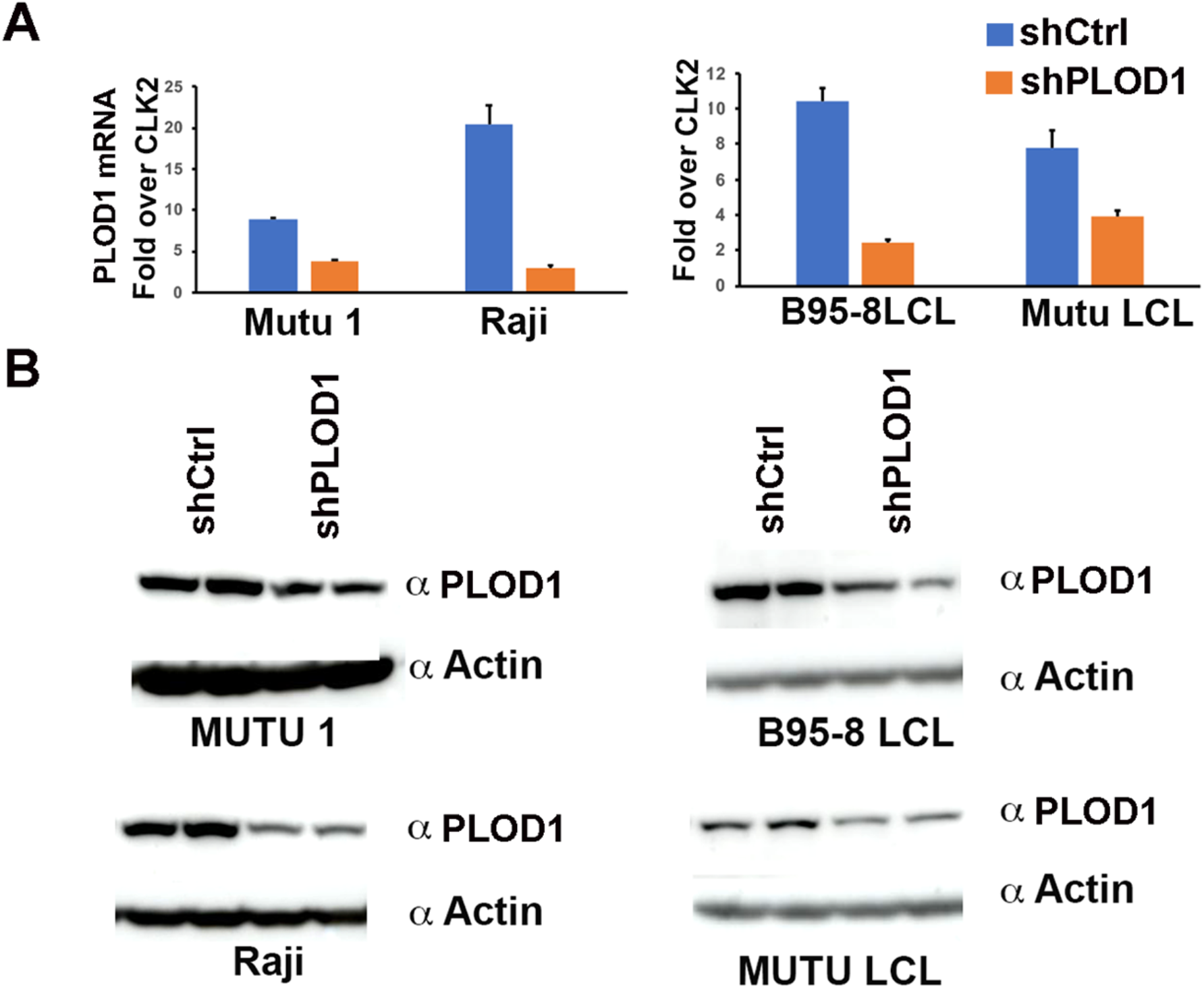
**A)** RT-qPCR analysis of PLOD1 mRNA in MutuI or Raji BL cells, or B95-8 or Mutu LCLs transduced with shCtrl or shPLOD1. **B**) Western blot of cells treated as described for panel A, and probed with antibody to PLOD1 (top panel) or Actin (lower panel). Each lane represents a biological replicate.

**Supplementary Figure S3.**
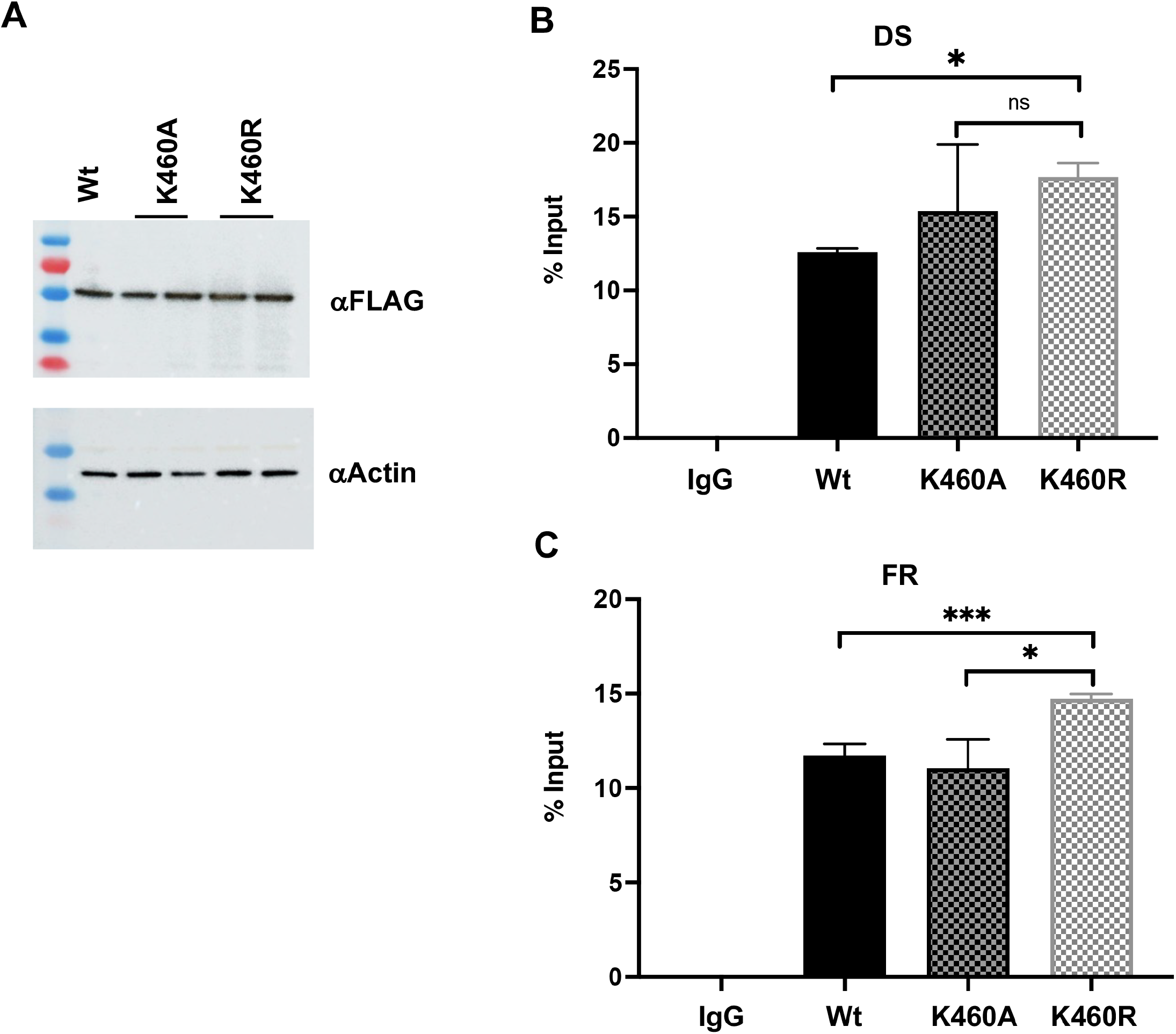
Mutations in K460 do not disrupt EBNA1 oriP binding in vivo. **A)** Western blot for FLAG-EBNA1 and Actin in HEK293T cells transfected with *oriP* plasmids expressing FLAG-EBNA1 Wt, K460A, or K460R. **B and C)** ChIP assays for control IgG or FLAG-EBNA1 Wt, K460A, or K460R at *oriP* DS region (**B**) or FR region (**C**) for extracts shown in panel A. P-values determined by ordinary one-way ANOVA and Dunnett’s multiple comparison test ***<0.001, *<0.05

**Supplementary Figure S4.**
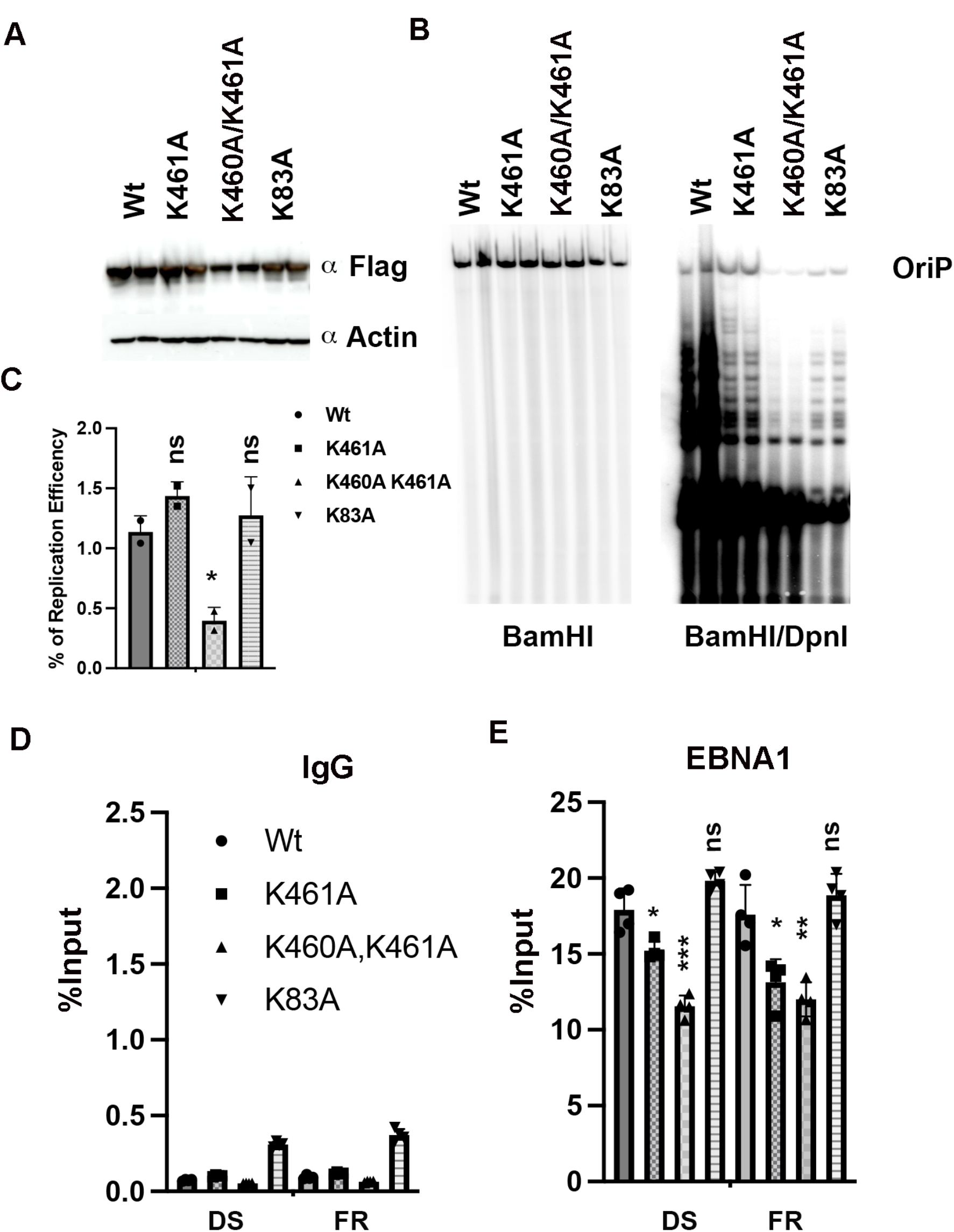
Mutations in K461 or K83A do not disrupt EBNA1 oriP binding in vivo. **A)** Western blot for FLAG-EBNA1 and Actin in HEK293T cells transfected with *oriP* plasmids expressing FLAG-EBNA1 Wt, K461A, K460A/K461A, or K83A. **B)** Southern blot of oriP replication for cells shown in panel A. **C)** Quantification of oriP replication shown in panel B. **D-E)** ChIP assay for control IgG (**E**) or FLAG-EBNA1 (**D**) or at *oriP* DNA for EBNA1 Wt, K461A, K460A/K461A, or K83A.. P-values determined by ordinary one-way ANOVA and Dunnett’s multiple comparison test ***<0.001, **<.01, *<0.05

